# Developmental molecular controls over arealization of descending cortical motor pathways

**DOI:** 10.1101/2023.06.29.546438

**Authors:** Philipp Abe, Adrien Lavalley, Ilaria Morassut, Esther Klingler, Antonio J. Santinha, Randall J. Platt, Denis Jabaudon

## Abstract

Layer 5 extratelencephalic (ET) neurons are a main class of neocortical projection neurons that predominate in the motor cortex and send their axon to the pons and spinal cord, and collaterals to the thalamus and multiple deep subcerebral structures^1–3^. Precise connectivity of ET neurons is critical for fine motor control; they are central to loss of function upon spinal cord injury and specifically degenerate in select neurodegenerative disorders^4, 5^. ET neurons consist of several types of cells with distinct laminar and areal locations, molecular identities, connectivities, and functions^6, 7^. Within layer 5 of the cortex, two cardinal subtypes of ET neurons have been identified: “ET_lower_” neurons, which express Slco2a1 and project to distal targets including the spinal cord, “ET_upper_“ neurons, which express Nprs1 or Hpgd and project more proximally to the pons and thalamus^6^. Despite their critical function, how these neuronal subtypes emerge during development and acquire their area-specific distributions remains unaddressed. Here, using combinations of anatomical labeling, MAPseq mapping^8^, and single-nucleus transcriptomics across developing cortical areas, we reveal that these two subtypes of ET neurons are present at birth along opposite antero-posterior cortical gradients. We first characterize area-specific developmental axonal dynamics of ET_lower_ and ET_upper_ neurons and find that the latter can emerge by pruning of subsets of ET_lower_ neurons. We next identify area- and ET neuron type-specific developmental transcriptional programs to identify key target genes *in vivo*. Finally, we reprogram ET neuron area-specific connectivity from motor to visual by postnatal *in vivo* combinatorial knockout of three key type-specific transcription factors. Together, these findings delineate the functional transcriptional programs controlling ET neuron diversity across cortical areas and provide a molecular blueprint to investigate and direct the developmental emergence of corticospinal motor control.

---

ET neurons, commonly referred to as “corticospinal” neurons, predominate in the motor cortex yet the molecular mechanisms controlling how they acquire their area-specific distributions and projections are still poorly understood^9–15^. To systematically assess the developmental area-specific dynamics of axonal projection of ET neurons, we injected a retrogradely-transported adeno-associated virus (AAV2 retro Nuc GFP) into the cervical spinal cord at postnatal day (P)3, P8, P12, P21, and P56, and characterized the areal distributions of retrogradely-labeled ET neurons at P70 — the presence of a nuclear localization signal allowed us to automate cell detection in clarified brains (**Fig. 1a**, **Methods**). This approach was also complemented and validated by using a non-viral labeling strategy (**Supplementary Fig. 1a-d**).

**Figure 1.**
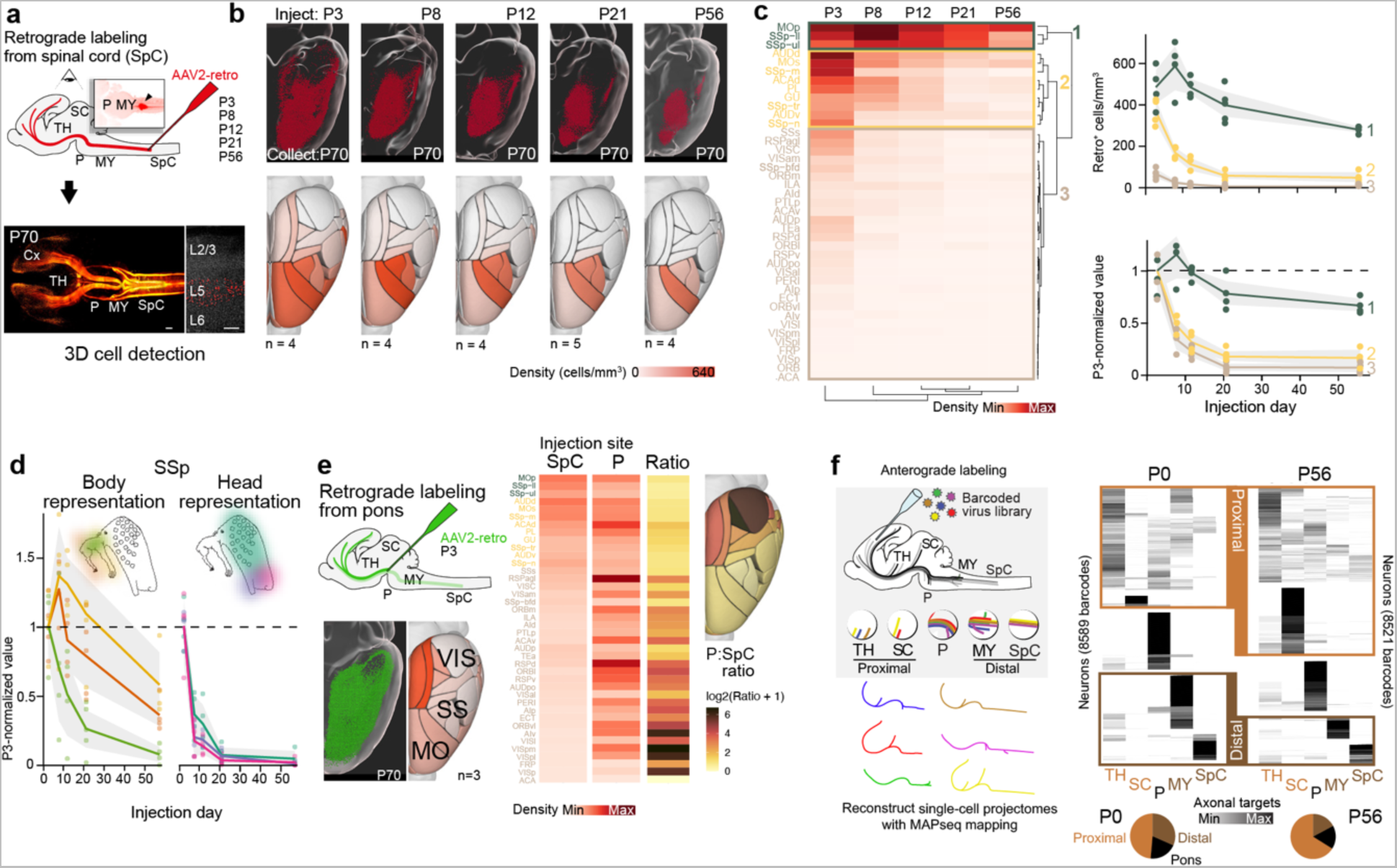
ET neurons have area-specific axonal dynamics during development. **a**, Top, Schematic illustration of the retrograde labeling from the spinal cord (SpC) at the indicated injection time points. Bottom left, example of a retrogradely-labeled clarified whole brain at P70; bottom right, retrogradely labeled nuclei visible across cortical areas in layer 5. **b,** Top-down view of retrogradely labeled cells (red) visualized in 3D-rendered brains and average cell density maps (cells/mm^3^) measured at P70 of mice injected at each respective time point (n=4 for P3, n=4 for P8, n=4 for P12, n=5 for P21, n=4 for P56). **c,** Left, heatmap and hierarchical clustering of average cell density detected in each cortical area (rows) at each injection timepoint (columns). Right, “pruning” curves of clustered cortical areas (cluster 1: green, cluster 2: yellow, cluster 3: brown) expressed as average cell density (top) and normalized values to average density at P3 (bottom). **d,** Average cell density normalized to P3 density for body representations (left) and head representations (right) of SSp. **e,** Top, Schematic illustration of the retrograde labeling from the pons (P) at P3. Bottom, example 3D reconstruction of detected cells at P70 (green) and corresponding cell density map. Right, Comparison of SpC and Pons (P) retrograde labeling experiments (injection: P3, analysis: P70) illustrated by cell density heatmap and brain map color coded by P:SpC ratio. Data from SpC is the same as in panel c. **f,** Left, schematic representation of the MAPseq mapping strategy. Right, Multiplex axonal projections of single ET neurons at P0+24 h and P56+24h. Heatmap representing projections clustered into proximal- (main targets in TH/SC) and distal- (MY/SpC) targeting. *Abbrevations:* Cx, cortex; MO, motor cortex; TH, Thalamus; SC, superior colliculus; P, pons; MY, medulla; SS, somatosensory cortex; SSp, primary somatosensory cortex; SpC, spinal cord; VIS, visual cortex. *Scale bars:* in **a** left 1 mm, right 200 µm. Abbreviations for the heatmap are according to the Allen Mouse Brain Atlas nomenclature (https://mouse.brain-map.org/static/atlas).

Three types of area-specific axon projection dynamics emerged from this analysis (**Fig. 1b,c**): first, anterior areas such as the motor cortex (MO) and primary somatosensory cortex (SSp), that establish robust projections early on and remain connected with the spinal cord; second, intermediate areas such as the auditory cortex and some regions of the SSp, that establish transient projections to the spinal cord that disappear during the first postnatal week; and third, posterior areas such as the primary visual cortex (VISp), with little or no projections to the spinal cord. At P3, the density of retrogradely-labeled neurons was high in regions that connected to the spinal cord, regardless of whether this connectivity was stable (as in anterior areas) or transient (as in intermediate areas); it was low or absent at all time points in posterior areas (**Fig. 1c**, top right). Together, these results confirm and extend previous findings^11, 15, 16^ by revealing early-onset area-specific differences in ET neuron axon projections that evolve during the first postnatal weeks.

Consistent with previous findings^11, 17, 18^, the postnatal reduction in projections from intermediate areas reflected axonal pruning rather than cell death. Indeed, cleaved Caspase3 staining did not reveal extensive or dynamic evidence of apoptosis (**Supplementary Fig. 1e**). Pruning rates were higher in transiently-projecting areas (i.e. intermediate areas) than stable ones (i.e. anterior areas) (**Fig. 1c**, bottom right), suggesting area-specific controls over ET axon stabilization. Remarkably, within SSp, pruning occurred in a somatotopic manner, with ET neurons located in head-representing areas undergoing rapid pruning, while ET neurons in body-representing areas pruned much less extensively (**Fig. 1d**). Retrograde labeling from the pons, which allows labeling of neurons with more proximal targets in addition to spinal cord-projecting ones, led to extensive labeling in more posterior regions than following spinal cord injections (**Fig. 1e**, left and middle); plotting pons-to-spinal cord projection ratios for each area revealed a posterior-high to anterior-low gradient, i.e. mirroring the one obtained with spinal cord injections. This suggests that areas with weak spinal cord projections instead project to more proximal targets (**Fig. 1e**, right). Confirming this possibility, anterograde labeling from MO, SSp and VISp using MAPseq mapping^8^ to identify proximal and distal ET targets in the thalamus, midbrain, pons, medulla and spinal cord confirmed axonal pruning, with superior colliculus (SC)-targeting axons forming the bulk of the newly developing proximal projections (**Fig. 1f**, **Supplementary Fig. 2**). Hence, while anterior areas project stably to the spinal cord, posterior areas instead project more proximally, while intermediate regions transiently project to the spinal cord before pruning their axons to more proximal targets. Together, these data suggest that early generation of ET neurons provides a ground state for future area-specific connectivity, after which cortical areas exhibit distinct patterns of axonal pruning and stabilization.

**Figure 2.**
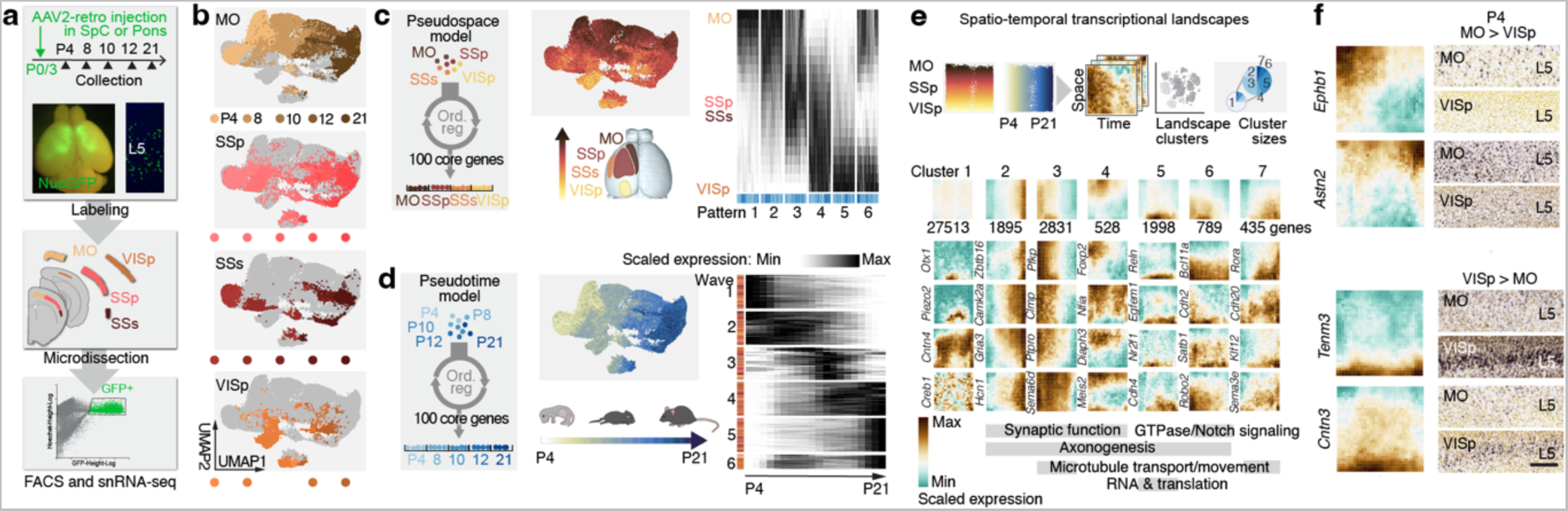
ET neurons have area-specific transcriptional dynamics during development. **a**, Schematic illustration of the experimental strategy. Photographs showing labeled cortical areas (left) and nuclei in L5 (right). **b,** UMAPs showing distribution of retrogradely-labelled ET neurons based on their transcriptional identity, color-coded by age of collection. **c,d,** Left, schematic illustration of the classification of ET neurons in cortical pseudospace (c) and pseudotime (d) using an ordinal regression models. Center, UMAP representation with color coded pseudospace (c) and pseudotime (d) prediction values. Right: Heatmap of expression dynamics (100 top core genes). **e,** Top, schematic illustration of the analysis pipeline for spatio-temporal transcriptional landscapes. Bottom, Mean expression landscapes for variable genes, single gene example landscapes and enriched GO terms. **f,** Examples of landscapes for genes enriched in the motor cortex (top) or the visual cortex (bottom). On the right of each landscape, the corresponding in situ hybridization data for L5 of motor and visual cortex are displayed (image source: www.brain-map.org). *Abbrevations:* MO, motor cortex; Ord.reg., ordinal regression; SSp, primary somatosensory cortex; SSs, secondary somatosensory cortex; SpC, spinal cord; snRNAseq, single-nucleus rNa sequencing; VISp, primary visual cortex. Scale bar: 300 µm.

To investigate the transcriptional programs that underlie these area-specific differences in ET neuron axonal properties, we first retrogradely labeled these cells from the spinal cord and the pons at P3 with AAV2-retro Nuc GFP and microdissected MO, SSp, the secondary somatosensory cortex (SSs) and VISp at P4, 8, 10, 12, and 21 before performing single-nucleus RNA sequencing of fluorescence-activated cell-sorted nuclei (**Fig. 2a**, **Supplementary Fig. 3a,b**). Analysis of cellular transcriptional identities was performed by Uniform Manifold Approximation and Projection (UMAP) dimensionality reduction. Two axes of transcriptional organization emerged from this analysis (**Fig. 2b**): (i) a spatial axis, which corresponded to the area identity of these cells across the tangential surface of the cortex, and (ii) a temporal axis, corresponding to the differentiation of ET neuron on successive postnatal days. These two cardinal features are thus the main source of transcriptional diversity of ET neurons in the developing neocortex. As previously described in adults, ET neurons clustered together except for a smaller cluster of *Chrna6*-type neurons^3^, whose spatial and temporal organization matched with that of the main cluster (**Supplementary Fig. 3c**).

**Figure 3.**
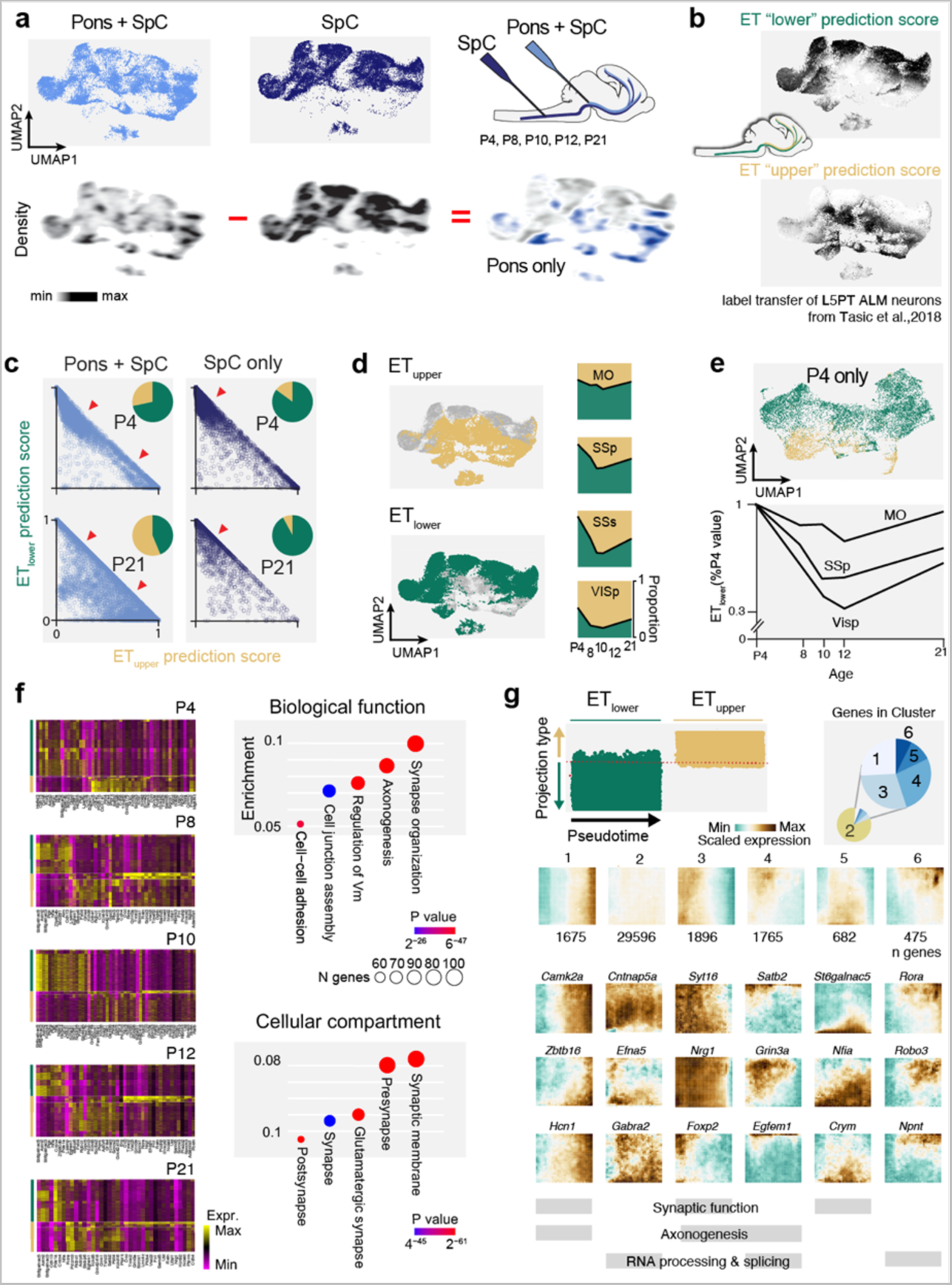
Developmental molecular parcellation of ET_upper_ and ET_iower_ neurons. **a**, UMAP (top) and corresponding density plot (bottom) for retrogradely-labeled neurons from the pons or spinal cord (schematic representation on the right). Density of pons-only projecting neurons is plotted by subtraction (highlighted in blue). **b,** Label transfer of ET_upper_ and ET_iower_ molecular identities from Tasic et al, 2018 onto the current dataset. ET_upper_ project proximally to the pons while ET_iower_ project to the spinal cord (schematic representation). **c,** Prediction values for Pons + SpC-projecting and SpC-only projecting neurons as ET_upper_ and ET_iower_ neurons (X and Y axes, respectively). Arrowheads highlight that SpC-only projecting are mostly predicted as ET_iower_, even at P4. **d,** Classification of ET neurons as ET_upper_ and ET_iower_ based on the label transfer (left). Proportion of each ET type across areas and postnatal age. **e,** Classification of ET neurons as ET_upper_ and ET_iower_ at P4 (top). While ET_iower_ predominate in MO, ET_upper_ progressively predominate in VISp (bottom). **f,** Left, Heatmaps of differentially expressed genes (DEGs) between ET_upper_ and ET_iower_-predicted neurons per age. Right, GO term analysis results for biological function (top) and cellular compartment (bottom) for DEGs. **g,** Top, Projection-type landscapes calculated for ET_upper_ and ET_iower_ across pseudotime. All genes were assigned to six landscapes cluster (pie chart) and mean landscape expression calculated on variable genes and representative landscapes are plotted. GO term enrichment is shown below. *Abbreviations:* MO, motor cortex; SSp, primary somatosensory cortex; SSs, secondary somatosensory cortex; SpC, spinal cord; VISp, primary visual cortex.

To address the spatial diversity of the molecular identity of ET neurons, we used an ordinal regression model in which single cells originating from the different cortical areas were ordered on a linear path based on their transcriptional identity. This created a pseudo-space alignment that outlined temporally-conserved transcriptional patterns driving the spatial diversity of ET neurons (**Fig. 2c**, **Supplementary Fig. 3d**). Spatially patterned transcripts included motor cortex-enriched genes such as *Crim1*^19^ and *Cdh12*^20^ (Patterns 1 and 2), visual cortex-enriched transcripts such as *Cntn6*^21^ and *Tenm3*^22^ (patterns 4 and 5), but also visual or motor cortex-excluded transcripts such as *Cntnap5a* and *Bcl11a*^23^ (patterns 3 and 4, respectively). Remarkably, ET neurons of the somatosensory cortex were found to be defined to a large extent by the presence of excluded genes (i.e. genes expressed in ET neurons of other areas but not the SSp, pattern 6), suggesting that thalamocortical input, which is particularly dense in SSp, acts on local cortical circuits to repress gene expression in ET neurons within this sensory area^24^.

We next addressed the temporal progression of ET neuron molecular identity using an ordinal regression model as described above, but this time based on the age of collected cells (**Fig. 2d**, **Supplementary Fig. 3e**). This approach identified successive transcriptional waves driving ET neuron differentiation across postnatal development, independently of area location. These genes included critical axon guidance genes, such as *Sox11*^25^ (wave 1) and *L1cam*^26^ (wave 2) as well as synaptic genes, like *Gabra4*^27^ (wave4) and *Gria2*^28^ (wave 5). Remarkably, there was minimal overlap between the core 100 genes defining the temporal differentiation of ET neurons and those encoding location (only 12/200 genes were shared) suggesting that the area-specific features of ET neurons reflect combinatorial interactions between largely independent area-specific ground states onto which shared differentiation programs are applied.

We next combined these two approaches to identify area- and differentiation stage-specific patterns of ET neuron gene expression. Each neuron was assigned a position score and a differentiation score based on the combined expression of corresponding core genes, and was then embedded within a two-dimensional matrix, allowing the generation of spatio-temporal transcriptional “landscapes” (**Fig. 2e**). To identify canonical features of gene expression, we performed a UMAP–based cluster analysis of all transcriptional maps (see **Methods**), revealing canonical clusters of genes with similar spatio-temporal expression dynamics. While most genes were expressed diffusely or with highly variable spatiotemporal specificities (cluster 1) others were temporally confined across cortical areas (i.e. late or early, clusters 2 and 3, respectively), or spatially restricted to frontal or occipital regions across differentiation (clusters 4-7). Ontological analysis revealed that genes within these clusters formed a coherent transcriptional program enriched in genes coding for proteins associated with axonal and synaptic function (**Fig. 2e**, bottom), consistent with their role in setting distinct axonal projection properties to differentiating ET neurons. *In situ* hybridization for select gene candidates using the Allen Brain Institute *in situ* dataset (https://developingmouse.brain-map.org/) confirmed such patterning (**Fig. 2f**) and differentially expressed gene analysis further highlighted transcriptional differences in time and space (**Supplementary Fig. 3f**). Hence, areal space and developmental stage are molecularly combinatorially encoded within developing ET neurons.

We next examined the molecular identity of neurons with proximal vs. distal axonal projections by comparing retrogradely-labeled neurons from pons or spinal cord. Since labeling from the pons not only labels genuine pons-projecting neurons but also *en passant* axons on their way to the spinal cord, we used a subtractive approach to identify the latter population. This strategy revealed a sharp molecular delineation of the molecular identities of spinal cord- and pons-projecting neurons (**Fig. 3a**). ET neurons can be divided into two main classes based on axonal reconstruction and single-cell RNA sequencing data (**Fig. 3b**)^6^: (1) ET_lower_ neurons (originally called L5PT ALM “lower” neurons, which express *Slco2a1* and project to distal targets such as the medulla and spinal cord, and (2) ET_upper_ neurons, which express *Hpgd* or *Nprs1* (originally called L5PT ALM “upper” neurons), that project more proximally to the pons, midbrain and thalamus (**Fig. 3b**). Taking advantage of this taxonomy linking projection target to molecular identity, we built a model predicting ET_upper_ vs ET_lower_ neuron identity in our dataset (**Supplementary Fig.4**, **Methods**). Consistent with this classification, spinal-cord projecting neurons were mostly predicted as ET_lower_ neurons (**Fig. 3c**), even at P4. ET_lower_ and ET_upper_ neurons are present across cortical areas, with ET_lower_ neurons predominating anteriorly and ET_upper_ neurons posteriorly (**Fig. 3d**). Remarkably, and consistent with a recent finding indicating an early distinction between brainstem- and spinal-cord projections in the sensorimotor cortex^10^, neurons with either ET_lower_ or ET_upper_ identities were already distinctly present at P4 (**Fig. 3d,e**), and then progressively refined in an area-specific manner: proportions of ET_lower_ and ET_upper_ ratios were relatively stable across time in anterior areas, while ET_upper_ identity progressively increased during development in more posterior areas (**Fig. 3d,e**). Together, these results reveal an early postnatal ground state of mostly ET_lower_ neuronal identity that is later refined to allow postnatal development of ET_upper_ identities along an antero-posterior gradient, such that this neuronal type predominates in posterior regions from P8 on.

**Figure 4.**
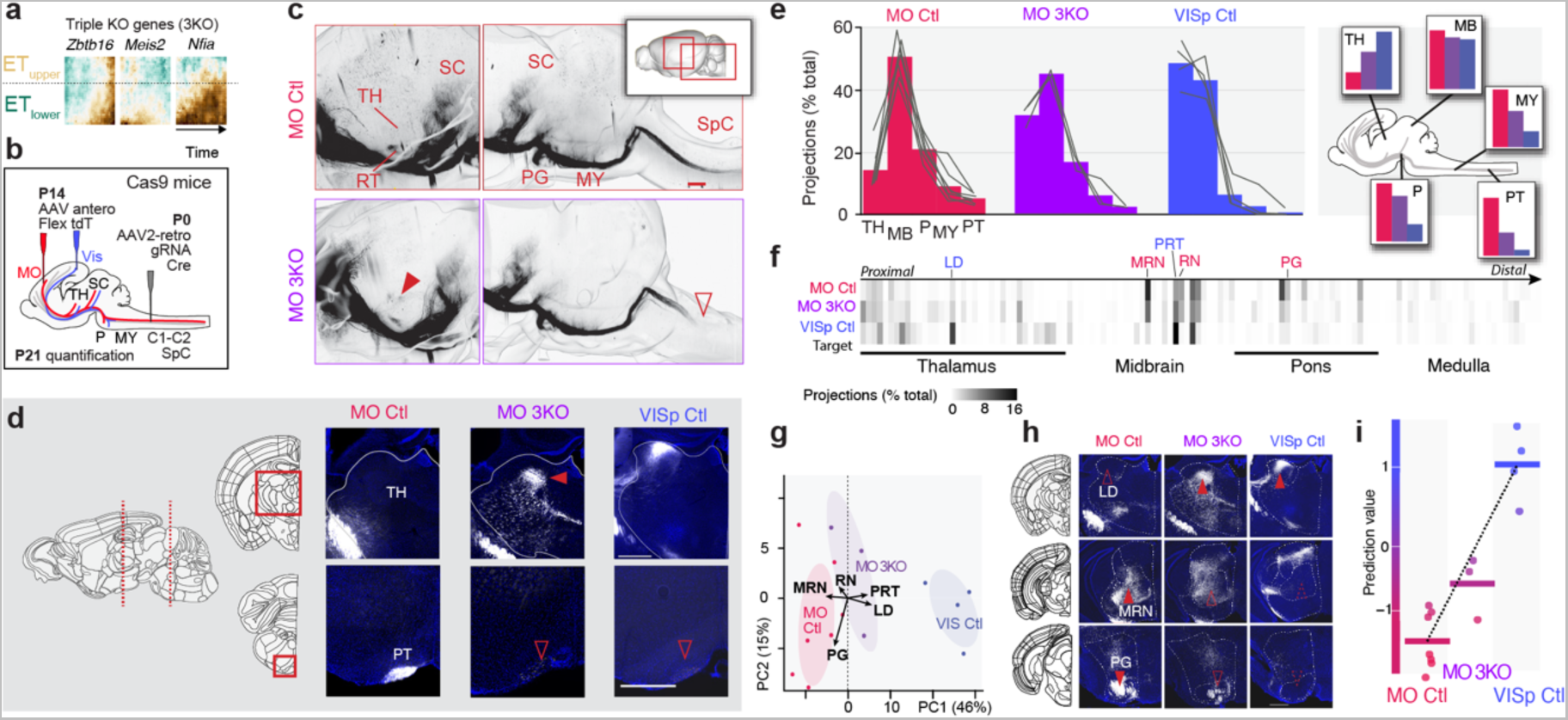
Genetic reprogramming of ET subtype connectivity. **a**, Projection-type landscapes for the three transcription factors targeted for CRISPR-mediated ablation (3KO). **b,** Experimental schematic for neuron projection-based genetic targeting. **c,** Top, representative images of 3D reconstructions after anterograde tracing from MO. **d.** Representative photomicrographs of sections showing axons (in white) in the thalamus (top) and pyramidal tract (bottom) following anterograde labeling from MO or VIS. **e**. Left, Percentage of projections for each analyzed target region for each condition. Right, the same values are displayed for each condition grouped by target region. **f,** Heatmap of percentage of projections (measured by axon-positive area in serial sections) for each condition ordered from proximal (left) to distal (right) targets. **g,** PCA analysis for relative projection to 131 subcortical brain regions (shown in f); each dot is one biological replicate (n=7 for MO Ctl; n=3 for 3KO; n=4 for VISp Ctl). **h,** Representative images of sections showing axons (in white) in thalamus (top panels), midbrain (middle panel) and pons (bottom panels). Structure selection is based on PCA eigenvectors in g. **i.** Scatter plot of prediction values for each condition, obtained from SVM machine learning model calculated on MO Ctl and VISp Ctl with the same input than the PCA. *Abbreviations:* Ctl, control; LD Lateral dorsal nucleus of thalamus; MRN Midbrain reticular nucleus; MY, medulla; MO, motor cortex; RN, red nucleus; P, pons; PG, Pontine grey; PRT, Pretectal region; PT, pyramidal tract; SSp, primary somatosensory cortex; SC, superior colliculus; SSs, secondary somatosensory cortex; SpC, spinal cord; tdT, td-tomato; TH, thalamus; TFmix, transcription factor mix; VIS, visual cortex. *Scale bars:* c, 1mm; d, h, 500 µm.

Genes distinguishing ET_lower_ and ET_upper_ neurons across postnatal development mostly encoded for proteins involved in axonal and synaptic function (**Fig. 3f**), consistent with the distinct axonal targets and functional role of these two cell types^6^. This included for example the cell adhesion protein *Cdh13*^29^ and the axon guidance molecule *Robo1*^30^, which are critically involved in axonal guidance. Using the same strategy as the one applied for developmental areal diversity of ET neurons (**Fig. 2e**), we established temporal landscapes for dynamically-expressed ET_lower_ and ET_upper_ genes (**Fig. 3g**). While most temporally-regulated genes were expressed with little cell-type specificity (clusters 1-3, representing 92% of all genes), a subset of transcripts were differentially expressed (clusters 4-6), suggesting sparse genetic encoding of ET identity. These genes included key neuron developmental molecules, such as *Satb2* (cluster4)^31^, *Nfia* (cluster5)^32^, *Rora*^33^ and *Robo3* (cluster6)^34^. These data identify thus exists distinct temporally-regulated genetic programs between ET_lower_ and ET_upper_.

The gradual appearance of ET_upper_ identity in the posterior cortical neurons suggests that there are transcriptional programs at play in ET_lower_ neurons (which are molecularly defined earlier; **Fig. 3d,e**) that drive the emergence of ET_upper_ neuron identity during development. To manipulate such programs, we sought for genes that would be expressed with high cell-type specificity in ET_lower_ neurons early on. Given the critical role of transcription factors in orchestrating developmental programs of gene expression, we focused our investigation on such transcripts and identified three such genes (**Fig. 4a**): *Zbtb16*, a zinc finger transcription factor involved in deep layer neuron production^35^, *Meis2*, a homeobox protein which influences neuronal fate in the striatum^36^ and retina^37^, and *Nfia*, a nuclear transcription factor which controls several aspects cortical development^38^, such as progenitor differentiation^32^ and corpus callosum formation^39^. We reasoned that loss-of-function of these genes in ET_lower_ neurons, in isolation or in combination, could cause the acquisition of ET_upper_ neuron-type projections. To specifically target ET_lower_ neurons, we retrogradely labeled these neurons at P0.5 from the spinal cord using an AAV2-retro viral vector containing a guide RNA-Cre-GFP construct targeted against each of the three transcription factors of interest (**Supplementary Fig. 5a**). We performed these experiments in Cre-dependent Cas9-expressing transgenic mice^40^ such that expression of the construct will lead to genomic ablation of the target gene in L5 ET neurons (**Fig. 4b**). Finally, to specifically follow the fate of reprogrammed ET_lower_ neurons, we specifically anterogradely labeled construct-containing neurons from MO at P14 by using a Cre-dependent tdTomato-expressing AAV vector (**Fig. 4b**).

Assessment of axonal projections in 3D clarified brains and classical 2D sections revealed that while single loss of each of these transcription factors (TF) had limited or no effect on axonal projections (**Supplementary Fig. 5c-f**), ablation of these TF together led to a reassignment in axonal targets (**Fig. 4c-i**). In contrast to control ET_lower_ neurons, which strongly projected to the spinal cord and largely bypass the thalamus, simultaneous loss of TF expression led to the acquisition of ET_upper_ neuron-type projections, with the generation of thalamic collaterals (**Fig. 4c,d** full red arrowhead) and reduction of distal projections (**Fig. 4c,d** empty red arrowhead). Remarkably, this projection pattern resembled that found in visual cortical ET neurons, where ET_upper_ neurons predominate and where more proximal, thalamic projections are found (**Fig. 4e**). Further, unbiased analysis of all subcortical targets revealed a congruent reassignment in key motor- and visual-specific projections. This includes for example loss of projections to the pontine gray matter (PG, normally a key motor relay center for corticopontine projections to the cranial nerves and cerebellum) and acquisition of projections to latero-dorsal nucleus thalamus (LD, normally a target of the visual cortex involved in spatial memory^41^) (**Fig. 4f-h**). Consistent with a motor-to-visual shift in axonal projections, prediction of projection identity using motor and visual projections as a training set revealed a motor-to-visual shift in the multiplex projection identity of motor neurons following reprogramming (**Fig. 4i**).

In summary, our results provide insights into the development of the two main types of ET neurons. Both ET_upper_ and ET_lower_ neurons are present early postnally, as also suggested by recent anatomical data^10^, but ET_lower_ neurons initially predominate, particularly in anterior cortical areas, while ET_upper_ proportion progressively rises in posterior regions through dedicated developmental transcriptional programs. Hence, there are two subtypes of ET_upper_ neurons - one with a ground state ET_upper_ identity, and one that emerges from the maturation of ET_lower_ neurons. Spatially restricted environmental signals, such as those conveyed by thalamocortical afferents, may play a role in this identity shift, including through master fate regulators such as *Zbtb16*, *Meis2*, and *Nfia*. Remarkably, each of these three transcription factors are involved in other aspects of the developmental biology of neurons, including cell migration and division. This highlights the inherent modularity of cellular molecular controls, which enable multiple functions in diverse biological contexts. Also, the requirement for loss of all three genes to reassign axonal projections supports the presence of multiple parallel molecular controls over axonal extension, stabilization, and target specificity.

Spinal-cord projecting neurons are initially present across the neocortex but then are predominant in the motor cortex. Instead of directly growing towards their final target, many neurons thus refine their transcriptional identity to prune to specific, more proximal postsynaptic partners. Extrasomatic regulation, such as axonal translation and protein modifications, which play a critical role in axon guidance, pruning and stabilization^42, 43^, might leverage these ground-state soma-derived axonal projection programs to allow for fine target specificity and circuit assembly. While the current study focuses on the molecular and anatomical delineation of ET neurons across cortical areas, similar mechanisms are likely at play to regulate projections to distinct spinal cord segments and brainstem vs. spinal cord projections within sensorimotor cortex^10, 19, 44–46^. Despite the potential metabolic drawbacks of “exuberant” growth followed by area-specific pruning, such axonal refinement may serve to allow activity-dependent interactions between proximal and distal sites within transient circuits^47^. Such a process could optimize functional connectivity by allowing competition between cortical areas for subcortical targets, ultimately optimizing circuit layout for the generation and coordination of complex movements.

## ACKNOWLEDGEMENTS

We thank the Genomics Platform, Bioimaging and FACS Facility of the University of Geneva; Advanced Lightsheet Imaging Center of the Wyss Center Geneva; A. Benoit for technical assistance; J. Prados, N. Baumann and Q. LoGiudice for assistance with bioinformatics analyses, L. Frangeul for logisitic support; all members of the Jabaudon laboratory for their comments on the manuscript as well as members of the Tole laboratories for constructive exchanges during the project. The Jabaudon laboratory is supported by the Swiss National Science Foundation, the Carigest Foundation, the Société Académique de Genève FOREMANE Fund, and the European Research Council; PA was supported by EMBO (ALTF 136-2017) and IRP (P 185F) Postdoc Fellowship. RJP is supported by the Swiss National Science Foundation grants 31003A_175830, 205965, and 40B2-0_211573/1; ETH Zurich grant ETH-27 18-2; Botnar Research Centre for Child Health Multi-Investigator Project Grant; National Centres of Competence – Molecular Systems Engineering 51NF40-182895; Brain & Behavior Research Foundation grant 26606, Personalized Health and Related Technologies grants 2021-542 and PHRT-203, Swiss Cancer Research Foundation grant KFS-4863-08-2019, European Research Council grant 851021, Botnar Research Centre for Child Health Fast Track Call grant, EMBO grant 4217, Simons Foundation Autism Research Initiative grant 1010692, F. Hoffmann-La Roche Roche Access to Distinguished Scientist Program (ROADS-036), and the Fickel Family Fund. AS is supported by the Swiss National Science Foundation grant 31003A_175830 and National Centres of Competence – Molecular Systems Engineering 51NF40-182895.

## METHODS

### Mouse strains and husbandry

CD1 male and female mice used in retrograde tracing, MAPseq, and Retro-NucSeq experiments were obtained from Charles River Laboratory. Rosa26-LSL-Cas9 knockin line used for CRISPR experiments was obtained from Prof. Randall Platt and maintained at Charles River Laboratory, following the genotyping protocol described in Platt et al., 2014^40^.

The experimental procedures described were conducted in accordance with the Swiss laws and previously approved by the Geneva Cantonal Veterinary Authority. All mice were housed in the institutional animal facility under standard 12h:12h light:dark cycles with food and water *ad libitum*.

### Retrograde and anterograde injections

For retrograde labeling at P0.5 and P3, pups were injected with 200 nl retrograde tracer in either cerebral peduncle of basilar pons or in the cervical corticospinal tract (spinal cord level: C1/2 at P1, C3 at P3) by using ultrasound guidance (VisualSonics Vevo^TM^-770-230) as described previously^48^. Retrograde labeling between P8 and P56 was performed under stereotaxic guidance (Stoelting Co.). Under isoflurane anesthesia, the cervical spinal cord was surgically exposed and 220 nl tracer was injected into the cervical corticospinal tract at C3 (ML: 0.2 mm, DV: 0.5 mm; Nanoject II, Drummond # 3-000-026).

For anterograde tracing with AAVs or Sindbis barcoded-viral particles, mice were injected in MO, SSp, or VIS at indicated ages under stereotaxic guidance (Stoelting Co.). Under isoflurane anesthesia, an incision was made to expose the skull and Bregma or Lambda landmarks were visualized using a stereomicroscope. The skull was opened either by a needle (pups ≤P14) or a drill and a glass capillary (outer diameter ∼60 µm) attached to a Nanoject II delivered the viral suspension followed by pausing 7 min at the injection site and slow retraction. The skin was sutured, and mice quickly recovered after surgery on a heating pad.

A list of the coordinates used for each age & brain region, which tracers were used for each experiment, the number of replicates, and their collection timepoints are provided in Table below.

### Tissue clearing and whole brain imaging

Mice were transcardially perfused with 4% PFA/PBS and brains were postfixed with 4% PFA/PBS at 4°C overnight. Subsequently, brains were clarified following the CLARITY protocol ^49, 50^. Imaging of entire clarified brains were acquired using a light-sheet microscope (MesoSPIM ^51^) with Z stacks at 5 μm spacing and a zoom set at 0.8x resulting in an in-plane spatial resolution of 8 μm (2048x2048 pixels) (further specifications under https://www.campusbiotech.ch/en/facilities/imaging-and-microscopy). Tissue clearing and 3D imaging for retrograde tracing experiments was performed with viral tracers, Retrobeads and for CRISPR perturbation experiments (see Table below).

### 3D rendering, quantification and mapping to reference atlas

3D renderings and quantifications were performed using Imaris software (Oxford Instruments). AAV2-retro+ and Retrobead+ cells were detected using Spot detection algorithm followed by registration to the Allen mouse brain common coordinate framework (CCF) using MIRACL^52^and Fiji^53^.

### *In vivo* CRISPR-Cas9 knockout experiments

#### gRNA design and cloning

Individual gRNA sequences targeting genes of interest were designed with the online tool CRISPick^54^. Control safe-harbor targeting gRNAs were selected from a reference dataset previously established^55^. Single gene targeting constructs containing two different sgRNAs were cloned by PCR amplification of the U6 promoter with the 3’-overhang of the reverse primer, including the respective sgRNA and the tracer RNA^56^ sequence. The two PCR fragments were inserted between the MluI and XbaI sites of the plasmid backbone in one cloning step.

#### AAV production and purification

HEK293T cells at 80% confluence in a 15 cm dishes (HuberLab) were transiently transfected with 21 µg of an equal molar-ratio mix of the AAV genome, AAV serotype plasmid, and the adeno helper plasmid pAdDeltaF6 (Puresyn) using polyethyleneimine max (PEI Max). The culture medium was harvested and replaced with fresh medium without FBS 48h post-transfection. One day later, cells collected and centrifuged at 800 x g for 15 min. The cell pellet was resuspended in 12 mL AAV lysis buffer (50 mL of 1 M TRIS-HCl (pH 8.5), 58.44 g NaCl, 5 mL of 2 M MgCl2 in 1L) and flash frozen in liquid nitrogen. The supernatant was mixed with 5 x AAV precipitation buffer (400 g PEG 8000, 146.1 g NaCl in 1 L H2O), joined to the medium harvested previously, incubated for 2 h at 4 °C, and centrifuged at 3000 x g for 1 h at 4 °C. The pellet was subjected to three freeze-thaw cycles and incubated with SAN (Merck, 50 U per 15 cm dish) for 1 h at 37 C. Ultracentrifugation gradients were prepared by sequential pipetting of iodixanol solutions: 9 mL (15%), 6 mL (25%), 5 mL (40%), and 5 mL (54%). Gradients were centrifuged using a Beckman type 70 Ti rotor at 63,000 rpm for 2 h at 4 °C. The tubes were pierced at the bottom, 4 mL of gradient were allowed to pass through and discarded, and the next 3.5 mL (containing isolated AAV) were kept. The solution was diluted with PBS + 10% glycerol and centrifuged through a 15 mL Amicon 100 kDa MWCO filter unit (Amicon) at 1000 x g for 10 min. The resulting AAV solutions were aliquoted and flash-frozen in liquid nitrogen.

#### AAV quantification

Viral particles concentration was determined by ddPCR (BioRad). Briefly, 5 µL of isolated AAVs were treated with DNAse I (NEB, M0303S) before preparing ten-fold serial dilutions. Primers targeting the WPRE sequence (WPRE_fwd: CTTTCCCCCTCCCTATTG; WPRE_rev: CAACACCACGGAATTGTC; WPRE_probe: CACGGCGGAACTCATCG) were used for ddPCR reactions. Droplets, data collection, and analysis were done BioRad ddPCR apparatus according with the manufacture’s indications.

Of note, AAV constructs delivered gRNA and CRE and thus allowed conditional L5 ET perturbation and anterograde tracing via additional Cre-dependent AAV1-FLEX-tdTomato transduction.

#### Targeting ET_lower_

At P0.5, Rosa26-LSL-Cas9 mice of the same litter were injected into the C1/2 SpC with either control gRNAs or targeting gRNAs, and for the pups injected with a mixture of gRNAs-containing vectors targeting *Nfia*, *Zbtb16* and *Meis2* (called 3KO), the vectors were mixed with a 1:1:1 ratio (n= 7 for control gRNAs, n= 4 for *Nfia*, n=4 for *Zbtb16*, n=4 for *Meis2*, n= 3 for 3KO). Cre-dependent anterograde tracing on the same animals was performed from the motor cortex, both brain and spinal cord were collected for histological analysis at P21, as described in the dedicated methods sections.

#### Targeting ET_lower_ + ET_upper_

At P0.5, Rosa26-LSL-Cas9 mice were injected into the cerebral peduncle at the basilar pons to transduce both L5 PT subtypes with control gRNA-Cre expression constructs. At P14, 110 nl of Cre-dependent anterograde tracer was injected in visual/retrosplenial cortex (called Ctrl VIS), and brain and spinal cord were collected for histological analysis at P21.

### Immunohistochemistry

Mice were perfused with 4% PFA/PBS and brains were first fixed overnight in 4% PFA/PBS at 4 °C, and then cryoprotected at 4°C in 20% sucrose in PBS until sinking. The brains were then embedded in OCT and preserved at –80°C until cryosectioning at 60 µm (Leica CM3050). For free-floating immunostaining, sections were first washed in PBS with 0,1% Triton X and then incubated for 2 h at room temperature in blocking–permeabilizing solution containing 3% bovine serum albumin and 0.3% triton X-100 in PBS, and subsequently incubated for 2 days with cleaved caspase 3 (1:200, Cell signaling 9661, LOT38) primary antibody at 4 °C. Sections were then rinsed 3 times in PBS with 0.1% Triton X and incubated with the corresponding Alexa-conjugated secondary antibodies (1:500; Invitrogen) for 2 h at room temperature. Nuclei were stained with Hoechst/PBS for 15 min. Native fluorescence of viral traced neurons and axons were imaged (e. g. tdTomato in CRISPR experiments). Images were acquired with Zeiss Axioscan.Z1 slide scanner, equipped with a 10× NA 0.45 Plan Apochromat objective, and a Hamamatsu Orca Flash 4 camera.

### Tissue collection and sequencing Retro-NucSeq

At indicated postnatal ages, mice were sacrificed and 400 µm-thick acute coronal brain slices were cut in artificial cerebrospinal fluid (ACSF, in mM: 87 NaCl, 25 NaHCO3, 2.5 KCl, 1.25 NaH2PO4, 0.5 CaCl2, 7 MgCl2, 5 glucose, 75 sucrose, saturated with 95% O2, 5% CO2, pH 7.4) using a Leica VT1200S vibratome. MO, SSp, SSs and VIS were micro-dissected and stored at –80°C until further processing. Of note, due to very low retrograde labeling from the SpC (Fig. 1), VISp samples were only collected from pons injected mice. All other samples were taken from both pons and SpC injections.

Nuclei were isolated using EZ Nuclei isolation kit. All steps were done on ice and in low binding tubes. To reduce technical batch effect, tissue of postnatal ages P4+P12 and P8+21 were pooled and separated bioinformatically. In brief, the tissue was resuspended in 2 mL of ice-cold extraction buffer (Nuclei EZ prep, Sigma NUC101), dounce-homogenized (KIMBLE Dounce tissue grinder, Sigma D8938), and incubated for 5 minutes in a total of 4 ml EZ buffer. The extracted nuclei were collected by centrifugation (500 x g at 4°C, 5 minutes) and supernatant removal followed by resuspension and incubation in EZ buffer for 5 min. After centrifugation, nuclear pellet was washed in 4 ml washing buffer containing 1% BSA in PBS, with 50U/ml of SUPERase-In (Thermo Fisher, cat# AM2696) and 50U/ml of RNasin (Promega, cat# N2611), centrifuged and nuclei were resuspended in 1 ml of washing buffer, filtered through a 30 μm strainer, stained with Hoechst (Invitrogen H3570, 1:500) for 5-10 minutes and processed for fluorescence-activated nuclei sorting (FANS) on Beckman Coulter MoFlo Astrios. 20k nuclei were sorted and 42 µl of nuclei suspension was used to load the 10x Genomics snRNA-seq preparation, according to the manufacturer’s instructions (10X Genomics Chromium 3′ Gene Expression Kit, cat 1000075, 1000073, 1000078). The cDNA libraries were quality controlled using a 2100 Bioanalyzer and TapeStation from Agilent, and sequenced using HiSeq 2500 sequencer. FASTQ files from sequencing were used as inputs to the 10X Genomics Cell Ranger pipeline (v3.0.2) with pre mRNA transcript detection of GRCm38 mouse genome.

### MAPseq

Tissue was collected and processed as described in Klingler et al, 2021^25^. In brief, 24 h after Sindbis viral transduction at indicated ages, injection sites and putative targets (Table, see below) were microdissected from 400 µm-thick acute slices and stored in Trizol reagent (Cat 10296-010, Thermo Fisher scientific) at –80°C. RNA extraction, construction of barcode cDNA library and sequencing was performed as previously described in Klingler et al., 2021.

### Analyses

All bioinformatics were performed using R Statistical Software (v4.1.1; R Core Team 2021) and Bioconductor packages as described below. Graphs and heatmaps were plotted with *ggplot2* and *complexHeatmap*^57^ packages.

### 3D retrograde tracing data

The output of *Imaris* spot detection and *MIRACL* brain registration was combined resulting in assignment of cell counts to cortical areas. Cell density was calculated by dividing raw counts by structure volume derived from mouse cell atlas ^58^.

Fig. 1b, 1e, Supplementary Fig. 1: Brain heatmaps were generated by *Cocoframer* R package (https://github.com/AllenInstitute/cocoframer).

### Retro-seq data

Libraries were preprocessed using a minimum of 1000 genes and 0.25 % mitochondrial cutoff and DoubletFinder^59^ for doublet removal. The SCTransform^60^ workflow in Seurat (v.4.3.0^61^) was run separately on each batch. Off target cells were excluded by assigning cellular identity using a custom marker gene panel. UMAP (Uniform Manifold Approximation and Projection) representation, marker genes and pseudotime analysis^62^ were used to assign age in age-pooled samples followed by merging of all identified L5 ET nuclei in a common object. Dimensionality reduction and clustering were performed using PCA and Louvain, respectively, using the default parameters of the Seurat pipeline and a common set of highly variable gene between 10x index batches (intersect between 5000 single index and 5000 dual-indices resulting in 2900 common variable genes).

#### Pseudo-space and -time (Fig. 2)

Cells were aligned in time and cortical space based on Telley et al., 2019 and *bmrm* R package (Bundle Methods for Regularized Risk Minimization Package, author: Julien Prados, year: 2018, R package version 3.10). Briefly, regularized ordinal regression method was used to predict the cortical area (pseudospace of MO, SSp, SSs and VISp) and the cellular age (pseudotime with input P4, P8, P10, P12 and P21). The linear weight of the model is used to rank the genes according to their ability to predict each cell in time or space followed by a selection of 100 core genes (top 50 and bottom 50) for each model. The prediction scores are further defined by the linear combination of the core gene expression. The pseudospace was divided into three models (Supplementary Fig. 3d): MO-SSp-VIS (MSV), MO-SSp-SSs (MSS) and VISp-SSp-SSs (VSS). The MSV model was used to predict pseudospace of SSs derived cells and for plots in Fig. 2c and 2e.

To generate transcriptional dynamics (also called waves), cells were aligned based on their pseudospace or -time score and their gene expression along this axis was corrected using weighted expression in neighbor cells. This was performed separately for each postnatal age in pseudospace and for each area in pseudotime. All transcriptional patterns were normalized to the maximum value and only genes with dynamic changes were kept for downstream analysis (per wave: *maximum value – minimum values > 0.2*) including the clustering into six groups (waves) based on their distances along pseudospace or -time using *hclust* function of *stats* package (k = 6). The heatmap in Fig. 2 c, d shows 100 representatives of each “space and time wave”.

#### Transcriptional landscapes

Calculation and plotting were previously described^62^. Cells were organized on a 2D grid based on their age and area status score followed by linear adjustment so that the average predicted values for each cardinal feature was aligned on the relative knot of the grid using the *tanh* function. The 2D space divided into a 32 x 32 grid and gene expression at a given coordinate of the 2D space was further estimated as the average expression of its 100 nearest neighbors. All transcriptional landscapes were normalized to the maximum value. The transcriptional landscapes of the most variable genes (n = 2900) were further clustered by projecting genes onto a 2D UMAP space computed on 15 principal components and sent to knn clustering (K = 6). The average expression pattern was calculated for each cluster and the transcriptional maps of all remaining genes (n = 33189) were correlated to these six patterns. Selected examples in Fig. 2e, f correspond to genes directly assigned or most highly correlated to the corresponding cluster.

#### Prediction of L5 ET subtype identity and projections

Published L5 ET subtypes and Retro-seq labels^6, 7^ were transferred to developing L5 ET neurons by applying *FindTransferAnchors* and *TransferData* functions included in Seurat package ^63^ (Supplementary Fig. 4a, parameters: normalized.method= “SCT”, recompte.residuals=F; PCA dimensions 1 to 30). Label transfer was verified by marker gene expression (Supplementary Fig. 4). *TransferData* also provides prediction values on how close a queried cell is to a given identity. Prediction values for L5 ET subtypes were used to construct transcriptional landscapes (see above) with L5 ET subtype identity (here ETlower/*Slco2a* scores) as function of pseudotime (Fig. 3, Supplementary Fig. 4).

#### Differentially expressed genes

For Fig. 3 and Supplementary Fig. 3: Differentially expressed genes (DEG) were identified by Seurat function *FindAllMarkers* and *test.use=”MAST”* with default parameters^64^. Top genes were plotted using *DoHeatmap* function.

#### Gene ontology terms

For Fig. 2e and Fig. 3f,g, gene enrichment on biological processes was calculated and plotted using clusterProfiler package^65^. P-values were adjusted by the FDR method.

### MAPseq mapping of projection patterns

MAPseq barcodes were demultiplexed, filtered and transformed into a counting matrix using the sindbis_docker container (https://github.com/pradosj/docker_sindbis) and previously published parameters^25^. In brief, after running demultiplexing and filtering (*TH=10*) on each injection site separately, counting matrices were combined in one R object and quality controlled by excluding barcodes present in negative control samples (eye and olfactory bulb) and with a maximum outside of the injection site (“soma calling”^66^), respectively. Further, barcode counts were normalized by spike-in counts found in each target to avoid any experimental bias due to library preparation.

All barcodes (∼projection neurons) were clustered based on their relative projection distribution (also called projection strength=UMI counts in a given target divided by the total UMI counts) across all collected targets (Table, *clara* cluster function in *cluster* package) and putative L5 ET neurons were selected by having projections to the thalamus, brain stem and spinal cord. Of note, neurons projecting only to the thalamus were categorized as corticothalamic (CT) and excluded from further analysis. L5 ET projections were further clustered in five clusters (optimal cluster number was determined by *factoextra* package): two proximal (main targets in TH+P or SC), one pontine and two distal (main targets MY or SpC) (Supplementary Fig. 2).

### Quantification of axons in CRISPR perturbation experiments

Anterograde-tdTomato + axons were quantified using a pixel classifier in Qupath^67^ and registered to the Allen mouse brain CCF using the ImageJ plugin ABBA (https://biop.github.io/ijp-imagetoatlas/). Axons in ipsi- and controlateral hemispheres were summed followed by averaging all axons within a structure. Further, axon signals were expressed as relative distribution per animal (axon fraction=axon signals in structure/total axon signals per animal).

For PCA and Support vector machine learning (SVM), relative axon fraction in 131 subcortical targets were used as input. *Bmrm* package was used to train a SVM on MO-Ctl and VIS-Ctl samples and perturbations groups were deduced by the resulting model (high values towards VIS-Ctl and low values towards MO-Ctl (Fig. 4, Supplementary Fig. 5e,f)

In Fig. 4e, Supplementary Fig. 5d, perturbation experiments were statistically compared to MO-Ctl by using a beta regression model (*betareg* package^68^) followed by “S*imulateneous Tests for General Linear Hypotheses*” for repeated measurement correction (*glht* function, *multcomp* package^69^).

#### Virus and Tracers

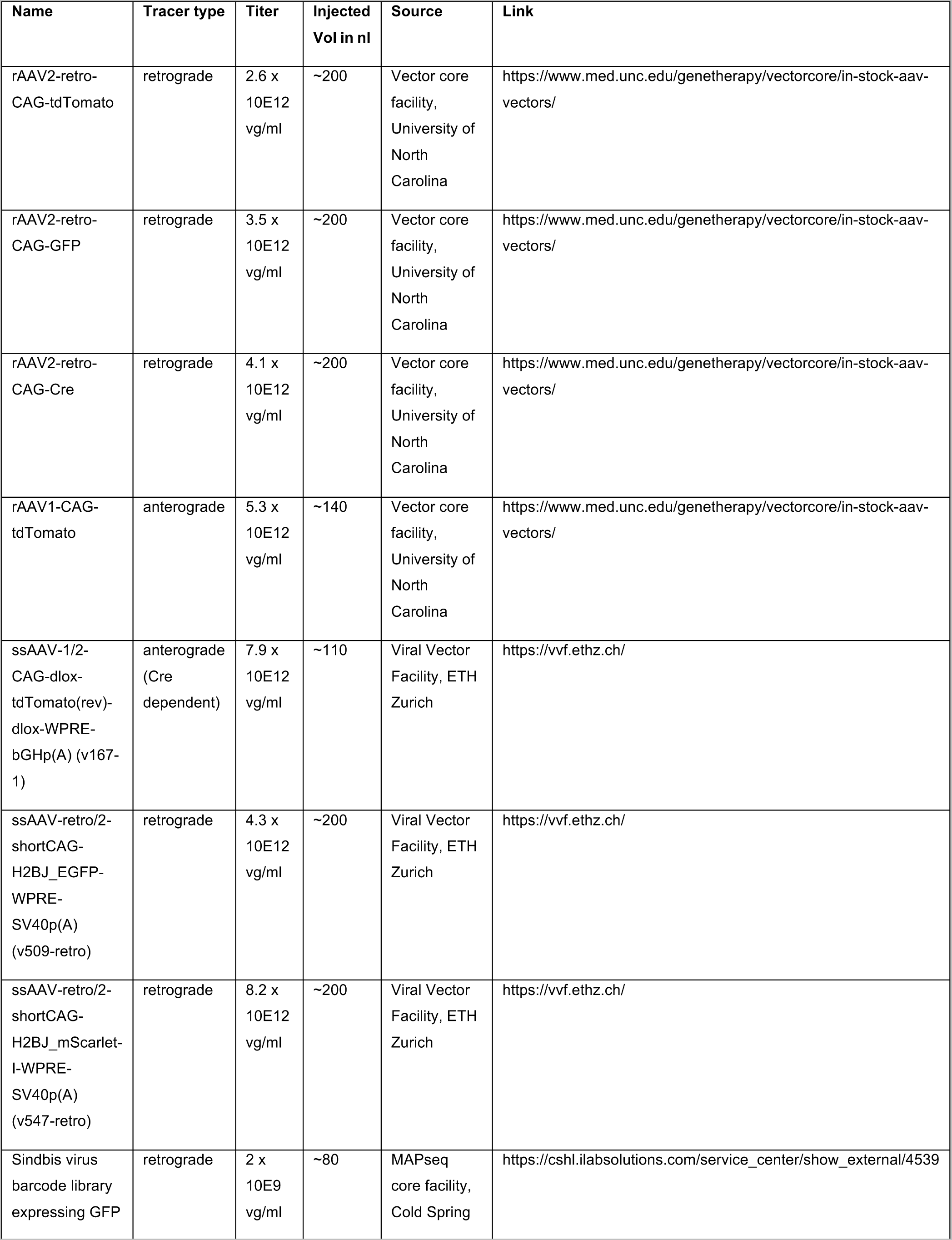
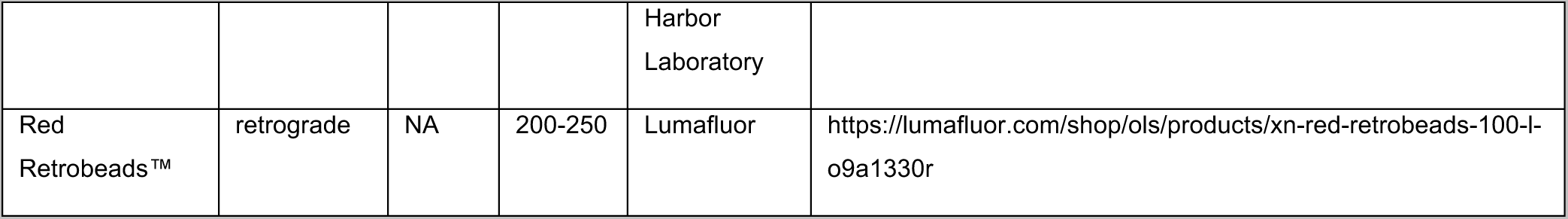

#### Coordinates

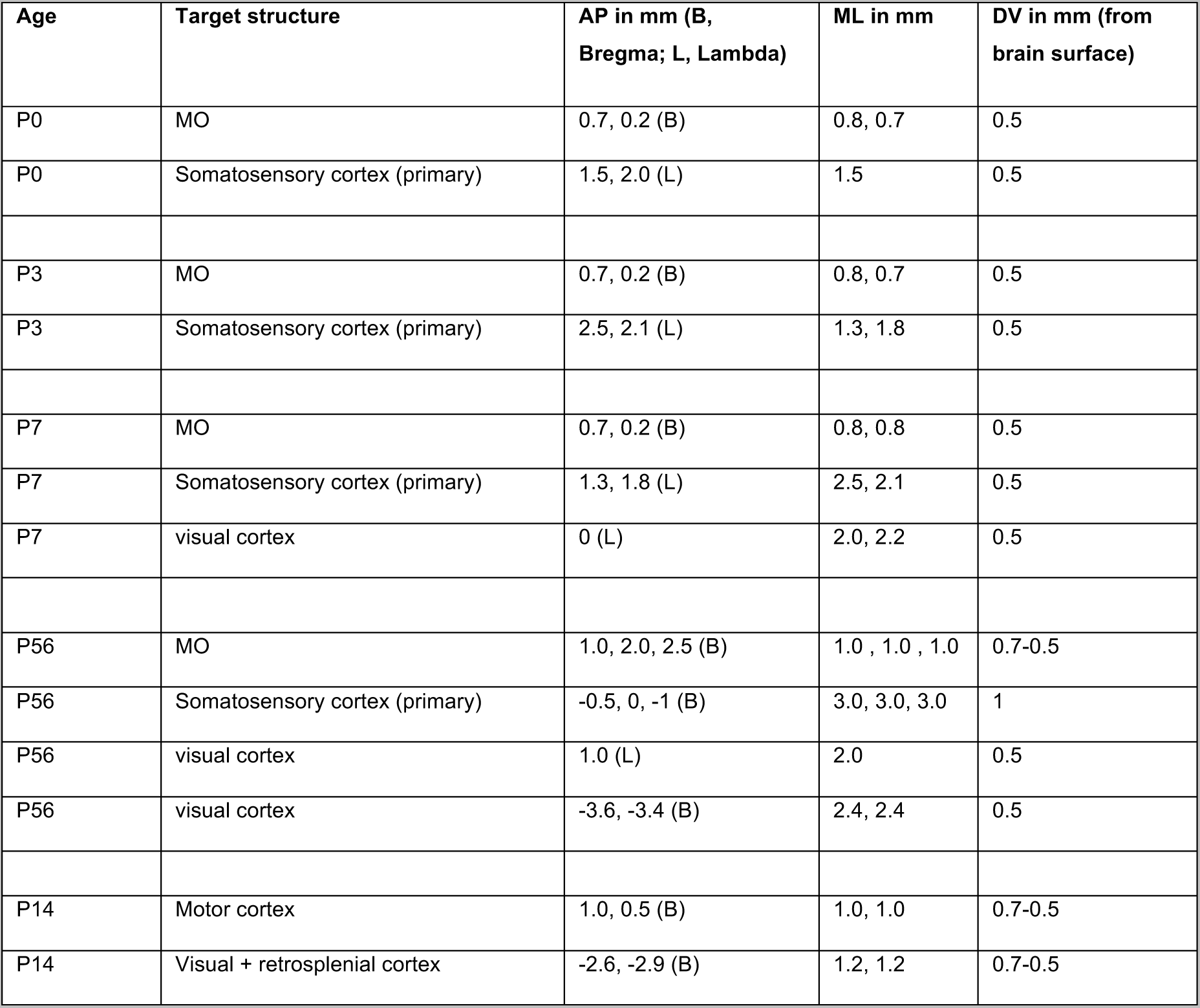

| Age | Target structure | AP in mm (B, Bregma; L, Lambda) | ML in mm | DV in mm (from brain surface) |
| --- | --- | --- | --- | --- |
| P0 | MO | 0.7, 0.2 (B) | 0.8, 0.7 | 0.5 |
| P0 | Somatosensory cortex (primary) | 1.5, 2.0 (L) | 1.5 | 0.5 |
| P3 | MO | 0.7, 0.2 (B) | 0.8, 0.7 | 0.5 |
| P3 | Somatosensory cortex (primary) | 2.5, 2.1 (L) | 1.3, 1.8 | 0.5 |
| P7 | MO | 0.7, 0.2 (B) | 0.8, 0.8 | 0.5 |
| P7 | Somatosensory cortex (primary) | 1.3, 1.8 (L) | 2.5, 2.1 | 0.5 |
| P7 | visual cortex | 0 (L) | 2.0, 2.2 | 0.5 |
| P56 | MO | 1.0, 2.0, 2.5 (B) | 1.0 , 1.0 , 1.0 | 0.7-0.5 |
| P56 | Somatosensory cortex (primary) | -0.5, 0, -1 (B) | 3.0, 3.0, 3.0 | 1 |
| P56 | visual cortex | 1.0 (L) | 2.0 | 0.5 |
| P56 | visual cortex | -3.6, -3.4 (B) | 2.4, 2.4 | 0.5 |
| P14 | Motor cortex | 1.0, 0.5 (B) | 1.0, 1.0 | 0.7-0.5 |
| P14 | Visual + retrosplenial cortex | -2.6, -2.9 (B) | 1.2, 1.2 | 0.7-0.5 |

#### MAPseq targets

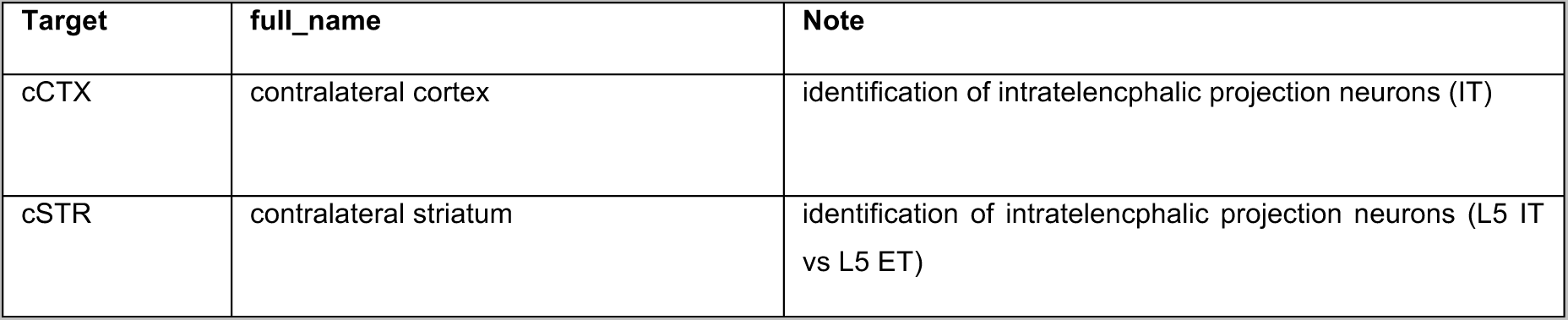
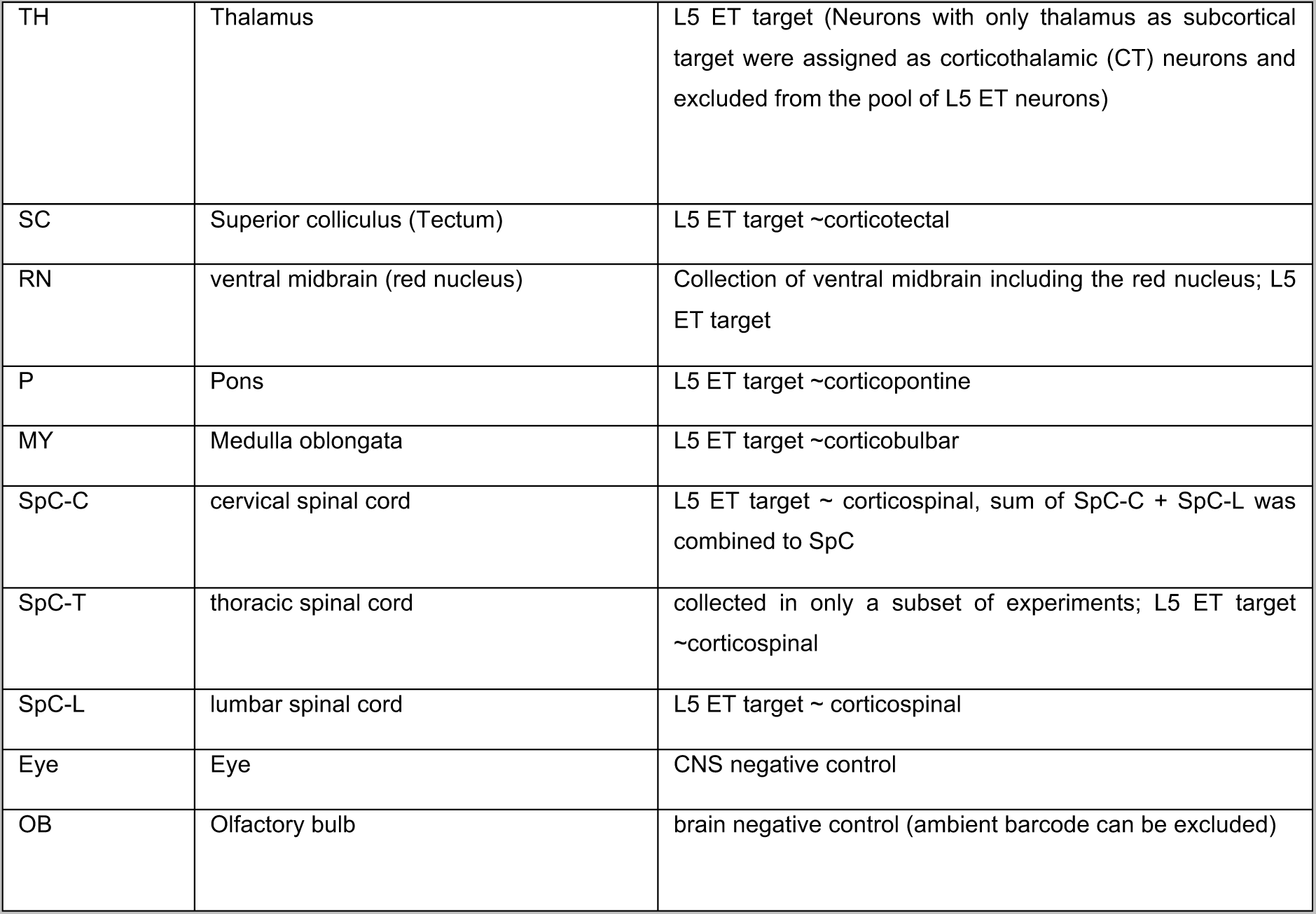

---

**Supplementary Figure 1.**
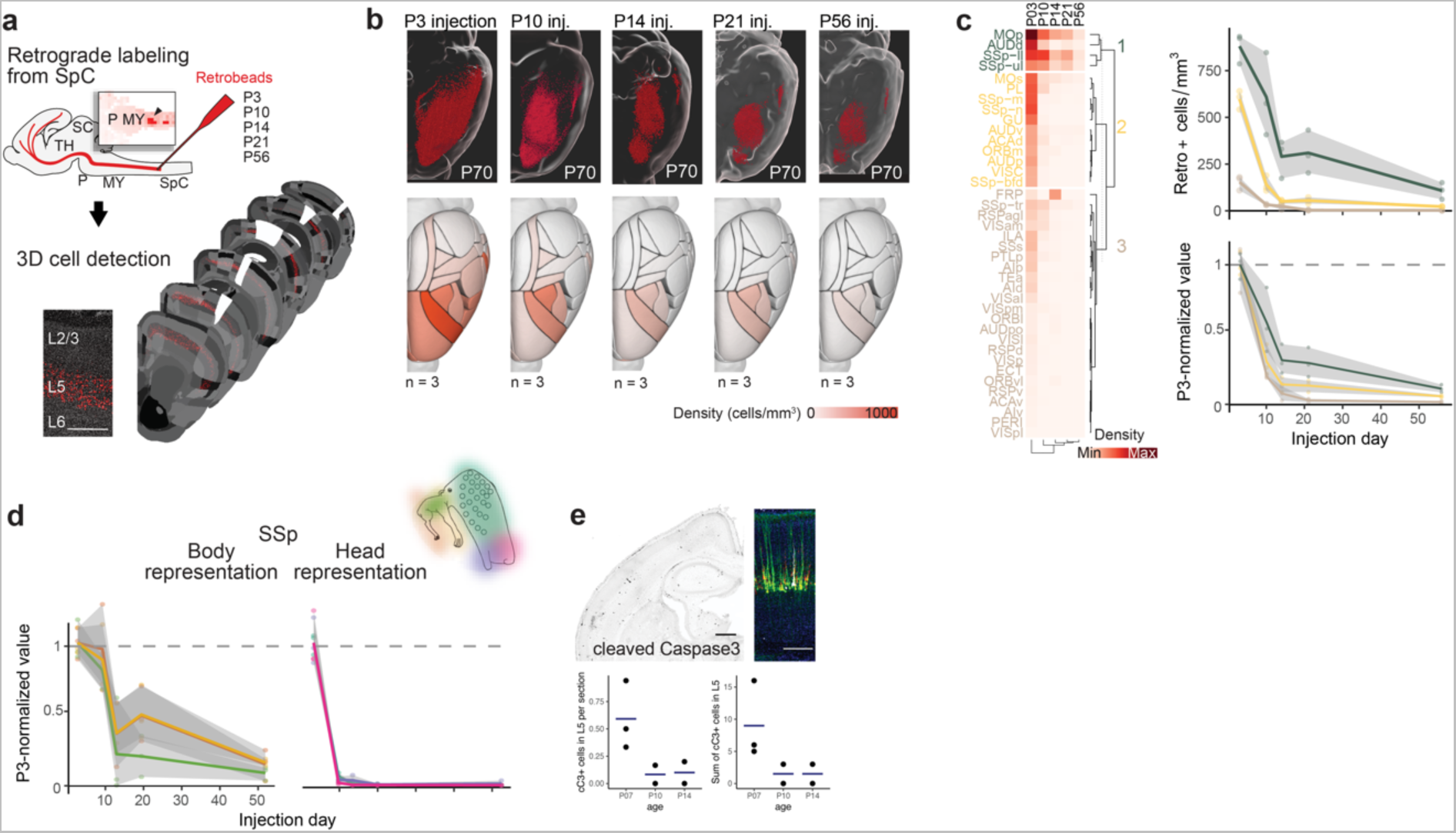
ET neurons have area-specific axonal dynamics during development. **a.** Top, Schematic illustration of the retrograde labeling procedure using retrograde labeling with fluorescent latex microspheres injected in the spinal cord (SpC). Bottom, At P70, coronal sections display retrogradely labeled cells detected across the cortex (left panel shows specific labeling of neurons in cortical layer 5). **b.** Top view of retrogradely labeled cells (red) visualized in 3D-rendered brains and average cell density maps (cells/mm^3^) measured at P70, of mice injected at each time point (n=3 for P3, n=3 for P10, n=3 for P14, n=3 for P21, n=3 for P56). **c.** Left, heatmap and hierarchical clustering of average cell density detected in each cortical area (rows) at each injection timepoint (columns). Right, “pruning” curves of clustered cortical areas (cluster 1: green, cluster 2: yellow, cluster 3: brown) expressed as average cell density (top) and normalized values to average density at P3 (bottom). **d.** Average cell density normalized to P3 density for body representations (left) and head representations (right) of SSp. **e.** Representative images (top) and quantifications (bottom) at P7, P10 and P14 of apoptotic cell marker cleaved Caspase 3 on L5 cells (Inset: P7 cortex with AAV2-retro-GFP (green), red Retrobeads and cleaved Caspase+ L5 neuron in white). *Abbrevations*: Cx, cortex; MO, motor cortex; TH, Thalamus; SC, superior colliculus; P, pons; MY, medulla; SS, somatosensory cortex; SSp, primary somatosensory cortex; SpC, spinal cord; VIS, visual cortex. *Scale bars*: in **a** left 1 mm, right 200 µm. Abbreviations for the heatmap are according to the Allen Mouse Brain Atlas nomenclature (https://mouse.brain-map.org/static/atlas).

**Supplementary Figure 2.**
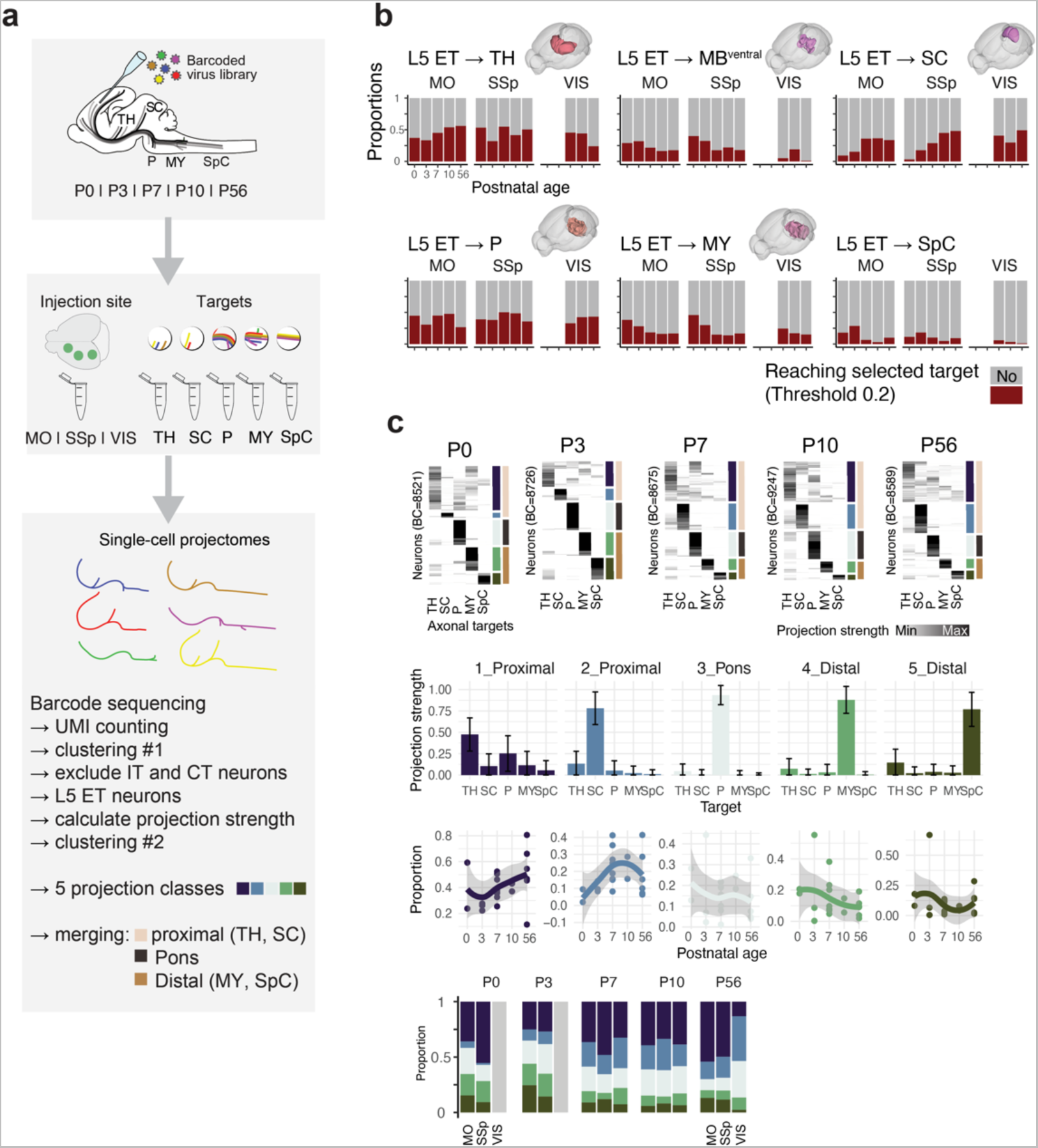
MAPseq mapping strategy. **a.** Schematic representation of the experimental and computational pipeline for MAPseq mapping of single-cell projections from motor, somatosensory and visual cortices. **b.** Proportions of projections detected in each cortical area for each of the individual targets (TH: thalamus, MBventral: ventral midbrain, SC: superior colliculus, P: pons, MY: medulla, SpC: spinal cord. Structures were considered as targets if projections to them represented > 20% of all projections). **c.** Top, Heatmap of projection strength at P0, P3, P7, P10 and P56. Each line represents one neuron/barcode, and each row represents the target analyzed. On the right of each graph, the first color-coding indicates the projection class, and the second one, the broader class of projection, each referred to the color-coding indicated in panel a. Middle, Bar graphs show the projection strength for each cluster based on detected barcodes in the five target regions and quantification of the proportions for each cluster of projection (2 proximal, 1 pontine, 2 distal) from P0 to P56 below. Bottom, bar plots of proportions for each projection cluster per timepoint.

**Supplementary Figure 3.**
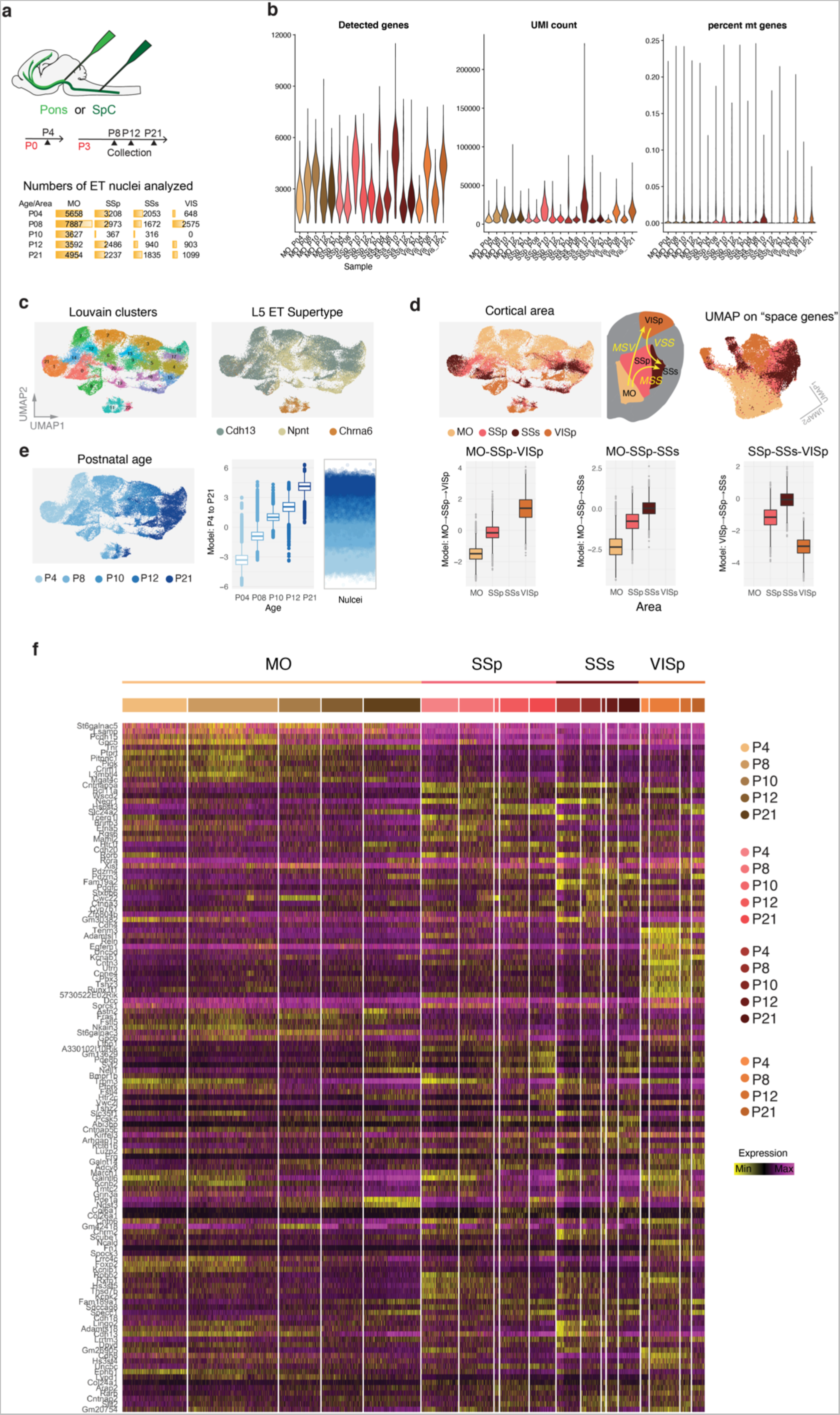
ET neurons have area-specific transcriptional dynamics during development. **a.** Top, Detailed schematic of Retro-NucSeq experiment referred to Figure 2. Bottom, exact numbers of nuclei included in the dataset per Area (columns) and timepoint of collection (rows) are reported. **b.** Violin plots for quality control parameters (number of genes per nucleus, UMI count and percentage of mitochondrial [mt] genes) per area and timepoint of collection are displayed. **c.** Left, UMAP representation of unbiased clustering. Right, UMAP representation shows the nuclei belonging to each supertype (Cdh13: 28477 nulcei, Npnt: 15825, Chrna6: 2408; Yao et al. 2021). **d.** Left, UMAP color-coded by cortical area. Right, Schematic summary of the three spatial ordinal regression models run on the dataset (MSV: motor-somatosensory-visual; MSS: motor-primary somatosensory-secondary somatosensory; VSS: visual-primary somatosensory-secondary somatosensory). ‘Space’ model derived genes (n=450 genes) were used to generate the UMAP representation on the right-end side. Note the similarity between genetic (UMAP) and anatomical space (scheme). Bottom, Boxplots for each of the three ordinal regression models representing the pseudospace alignment of nuclei derived from the different cortical areas. **e.** Top left, UMAP representation with color code for timepoint of collection. Top right, pseudo-time alignment of all nuclei for each collection timepoint represented as boxplot and scatter plot. **f**. Heatmap of differentially expressed genes (row) for each sequences nucleus (columns), grouped by area and timepoint of collection.

**Supplementary Figure 4.**
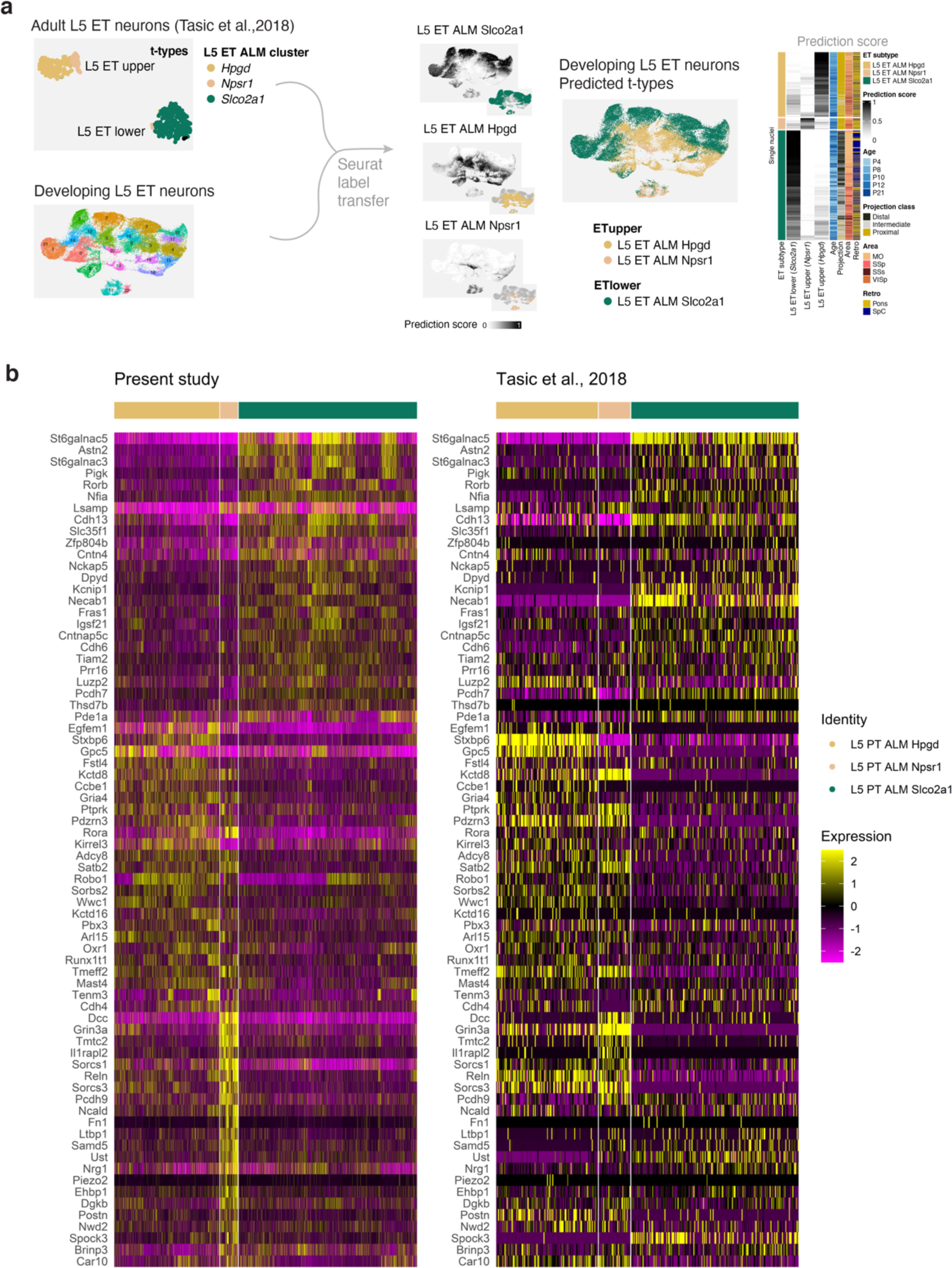
Identifying ET subtypes by using transcription-based label transfer. **a.** Left, Workflow of transferring labels from previously described transcriptional ET subtypes (Economo et al., 2018 and Tasic et al., 2018, dataset of adult L5ET ALM neurons) to present dataset of developing ET neurons. Middle, Feature plots showing the prediction score for the three subtype identities and color-coded UMAPs of the transferred subtype labels. Right, Heatmap of predictions scores for each nuclei (rows). **b.** Heatmaps of ET marker genes displayed in rows for each sequenced nucleus (columns) for the present study (left) and previously published dataset (Economo et al., 2018 and Tasic et al., 2018) (right). Note the similar subtype specific expression patterns in both datasets indicating a successful label transfer.

**Supplementary Figure 5.**
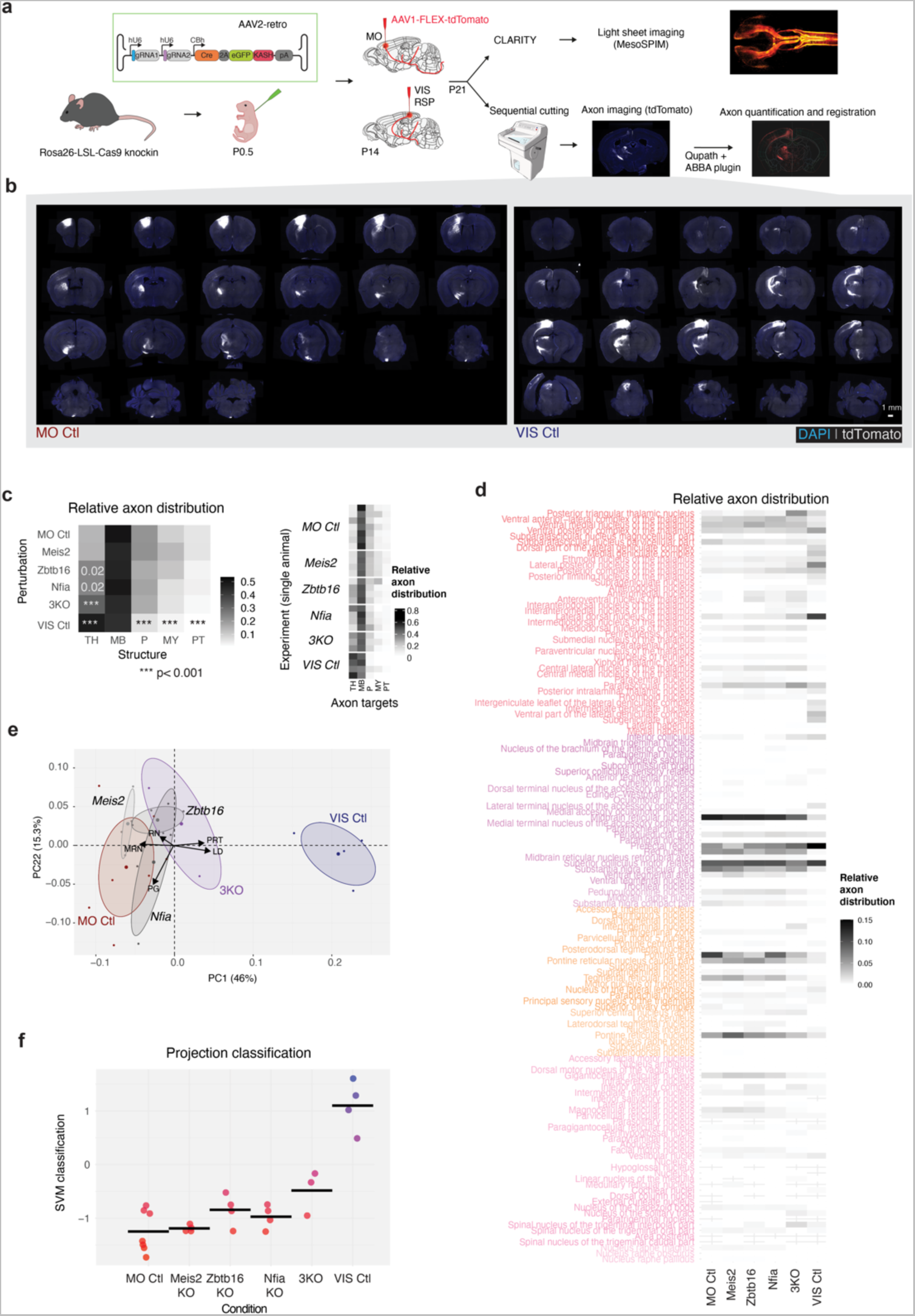
Reprogramming axonal connectivity with ET type-specific transcription factors. **a.** Experimental design **b.** Example of coronal sections showing the tdTomato+ axons (in white) obtained with anterograde tracing from the Motor cortex (left) and Visual cortex (right). **c.** Heatmap displaying the average relative axon distribution for each structure (columns) per condition. Significance test was calculated using beta regression and p-value below 0.001 are indicated with three asterisks. Values for single animals are displayed in the heatmap on the right. **d.** Heatmap representing the fraction of total projections detected for all the brain areas (rows) for each condition (columns). **e.** PCA of relative axon distribution across 131 targets per indicated condition. PCA values are represented per animal (small points) and as centroid (bigger points) for each condition. **f**. SVM classification score for each condition with dots representing biological replicates. SVM was trained on relative axon distribution of MO Ctl and VIS Ctl.

